# Widely used mutants of *eiger*, encoding the *Drosophila* Tumor Necrosis factor, carry additional mutations in the NimrodC1 phagocytosis receptor

**DOI:** 10.1101/2020.09.30.316257

**Authors:** Albana Kodra, Claire de la Cova, Abigail R. Gerhold, Laura A. Johnston

## Abstract

The process of apoptosis in epithelia involves activation of caspases, delamination of cells, and degradation of cellular components. Corpses and cellular debris are then rapidly cleared from the tissue by phagocytic blood cells. In studies of the *Drosophila* TNF, Eiger (Egr) and cell death in wing imaginal discs, the epithelial primordia of fly wings, we noticed that dying cells persisted longer in *egr^3^* mutant wing discs than in wild type discs, raising the possibility that their phagocytic engulfment by hemocytes was impaired. Further investigation revealed that lymph glands and circulating hemocytes from *egr^3^* mutant larvae were completely devoid of NimC1 staining, a marker of phagocytic hemocytes. Genome sequencing uncovered mutations in the *NimC1* coding region that are predicted to truncate the NimC1 protein before its transmembrane domain, and provide an explanation for the lack of NimC staining. The work that we report here demonstrates the presence of these *NimC1* mutations in the widely used *egr^3^* mutant, its sister allele, *egr^1^*, and its parental strain, *Regg1^GS9830^*. As the *egr^3^* and *egr^1^* alleles have been used in numerous studies of immunity and cell death, it may be advisable to re-evaluate their associated phenotypes.

## Results and Discussion

The *Drosophila* genome encodes a single TNF homolog, known as Eiger (Eda-like cell death trigger, Egr) (Igaki *et al*.; Moreno *et al*.; Narasimamurthy *et al.*). Egr is expressed in many different tissues and plays various roles in cellular processes such as the immune response, energy homeostasis, and JNK-dependent cell death. Since its identification, numerous studies on cell death and immunity have utilized the *egr^3^* and *egr^1^* alleles, which were generated by imprecise excisions of the *Regg1^GS9830^* P element and resulted in deletions of the first coding exon of the *egr* gene (Igaki *et al.*). Both *egr^3^* and *egr^1^* strains are homozygous viable and considered severe loss-of-function alleles.

Dead cells in *Drosophila* are commonly removed from tissues by phagocytic engulfment by plasmatocytes, the most abundant of the circulating hemocytes in the larva (Abrams *et al*.; Sonnenfeld and Jacobs; Franc *et al*.; Shklyar *et al.*). Plasmatocytes carry cell surface receptors for the recognition and rapid engulfment of bacteria, dead cells and cellular debris, such as Eater (Kocks *et al*.; Chung and Kocks), NimrodC1 (NimC1) (Kurucz *et al*.; Honti *et al.*) and Draper (Manaka *et al.*). A frequently used marker for plasmatocytes in *Drosophila* is positivity for NimC1, a transmembrane protein characterized by the presence of a special type of EGF repeat known as the NIM repeat, located immediately proximal to a conserved CCxGY motif (Somogyi *et al.*). The *NimC1* gene is part of a cluster of four *NimC (NimC1-4*) genes in the midst of several other related *Nimrod* genes at 34E on chromosome 2. Nimrod proteins contain 2–16 NIM repeats as well as additional conserved residues at their amino terminus. The Nimrod proteins, together with Eater and Draper, form a conserved superfamily of 12 proteins in *Drosophila*, and Nimrod proteins are also encoded in the *C. elegans* and mammalian genomes (Melcarne *et al.*). Loss of any Nimrod protein diminishes the capacity of hemocytes to fight microbes. For example, RNAi against *NimC1* has implicated it in bacterial phagocytosis (Kurucz *et al.*), while complete loss of *NimC1* demonstrated cooperativity between NimC1 and Eater in the recognition and phagocytosis of bacteria (Melcarne *et al.*).

Egr has also been reported to have a role in regulating phagocytosis of bacteria (Schneider 2007). In the course of studying the role of the Egr in cell competition, where apoptosis is non-cell autonomously induced, we found that dying cells appeared to persist in wing discs from *egr^3^* mutant larvae (**Fig. 1A-D**). That cell death was still induced in *egr^3^* mutant cells suggested that Egr is not required for the cells to die under these two conditions. However, because dead cells are typically cleared within 2-4 hours from wild-type wing imaginal disc epithelia (Milan *et al.*), the persistence of Cas-3 positive cells that we observed in *egr^3^* mutants suggested that loss of *egr* might impair corpse clearance. This prompted us to examine plasmatocytes in the lymph glands, the major larval hematopoietic organ, from WT and *egr^3^* mutant larvae. We immunostained the lymph glands from both genotypes with anti-NimC1 antibodies, a mixture of P1a and P1b antibodies that specifically recognizes the phagocytic plasmatocytes of the larva (Kurucz *et al.*). As a control, we also examined larvae that carried Hml-RFP, consisting of a hemocyte-specific enhancer/promoter from the *Hemolectin* gene fused to red fluorescent protein (RFP) that identifies larval hemocytes (Clark *et al*.)(**Fig. 2A**). NimC1 is expressed at high levels on the plasma membrane of numerous cells in the primary lymph gland lobes from WT controls, and anti-NimC1 staining overlapped with many Hml-RFP positive cells (**Fig. 2A, B**). Strikingly, however, no NimC1 positive cells were evident in lymph glands from *egr^3^* larvae (**Fig. 2C)**. As the *egr^3^* and *egr^1^* alleles were derived from the same parental strain (Igaki *et al.*), we also tested lymph glands from *egr^1^* larvae, and again found no detectable NimC1 expression (data not shown). To examine circulating plasmatocytes, we isolated hemocytes from larval hemolymph. Although NimC1 was readily observed in circulating hemocytes from *OregonR (OreR*) controls (**Fig. 2D**), we detected no NimC1-positive hemocytes in the hemolymph from *egr^3^/egr^1^* transheterozygous larvae, or from *egr^31^* larvae (**Fig. 2E-F)**. *egr^31^* is a precise excision of the *regg1^GS9830^* P-element present in the parental strain. Thus all of the *egr* alleles derived from the *regg1^GS983^* strain lacked circulating and lymph gland resident plasmatocytes that expressed NimC1.

**Figure 1.**
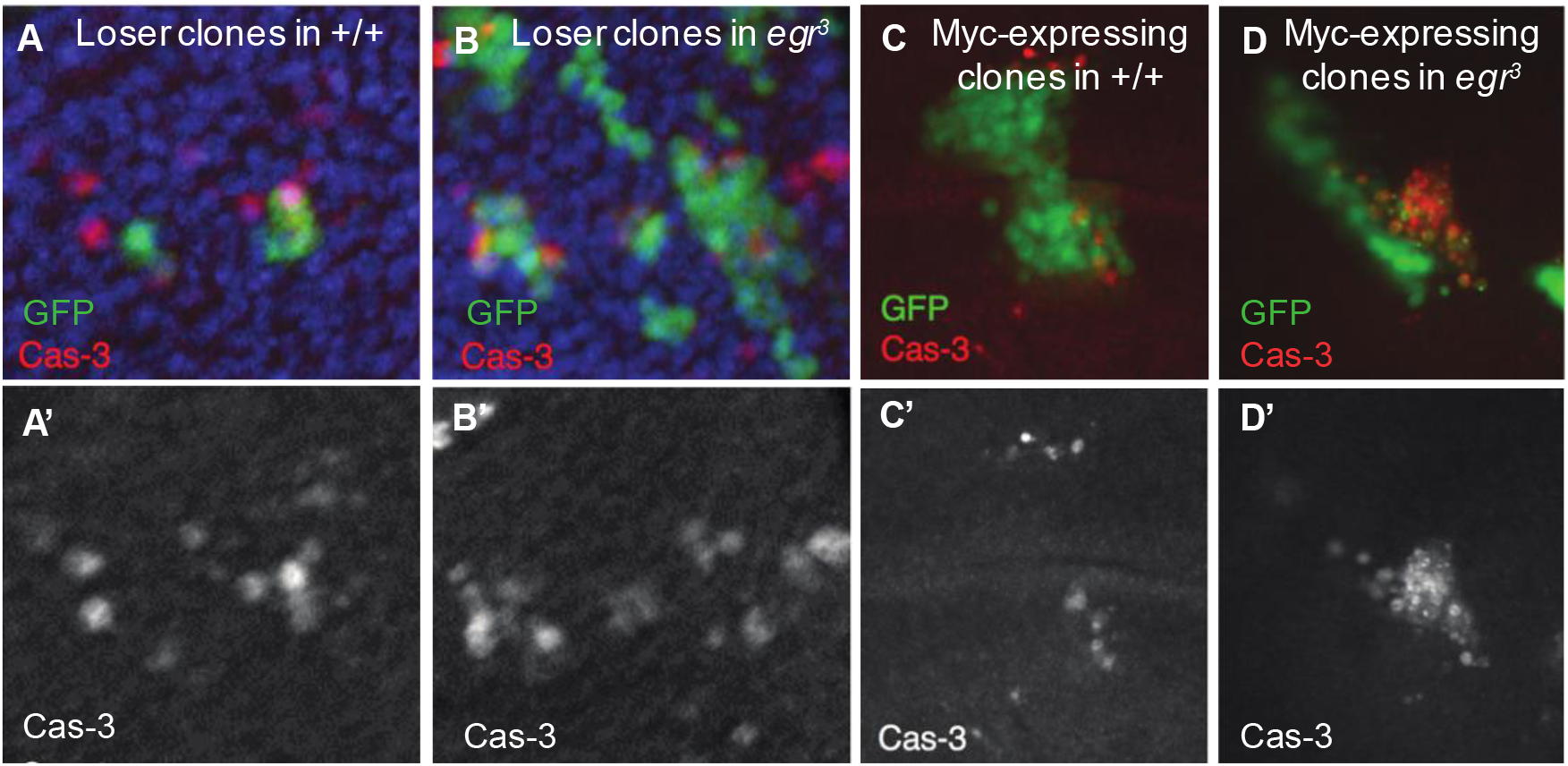
Dying cells may be cleared less efficiently in *egr^3^* mutants. A. Clones of loser cells in a wildtype wing disc, expressing GFP and cleaved caspase 3 (Cas-3); Cas-3 channel is shown in A’. B. Loser clones, expressing GFP and Cas-3, in a *egr^3^* mutant wing disc; Cas-3 channel is shown in B’. Clones in A and B were examined 24 hrs after clone induction. C. Myc-expressing clone, marked by expression of GFP, in a wildtype wing disc. Cas-3 positive cells are shown in red. C’ shows Cas-3 as a single channel. D. Myc-expressing clone, marked by expression of GFP, in an *egr^3^* mutant wing disc. Cas-3 positive cells are shown in red. D’ shows Cas-3 as a single channel. Clones in C and D were examined 48 hrs after clone induction.

**Figure 2.**
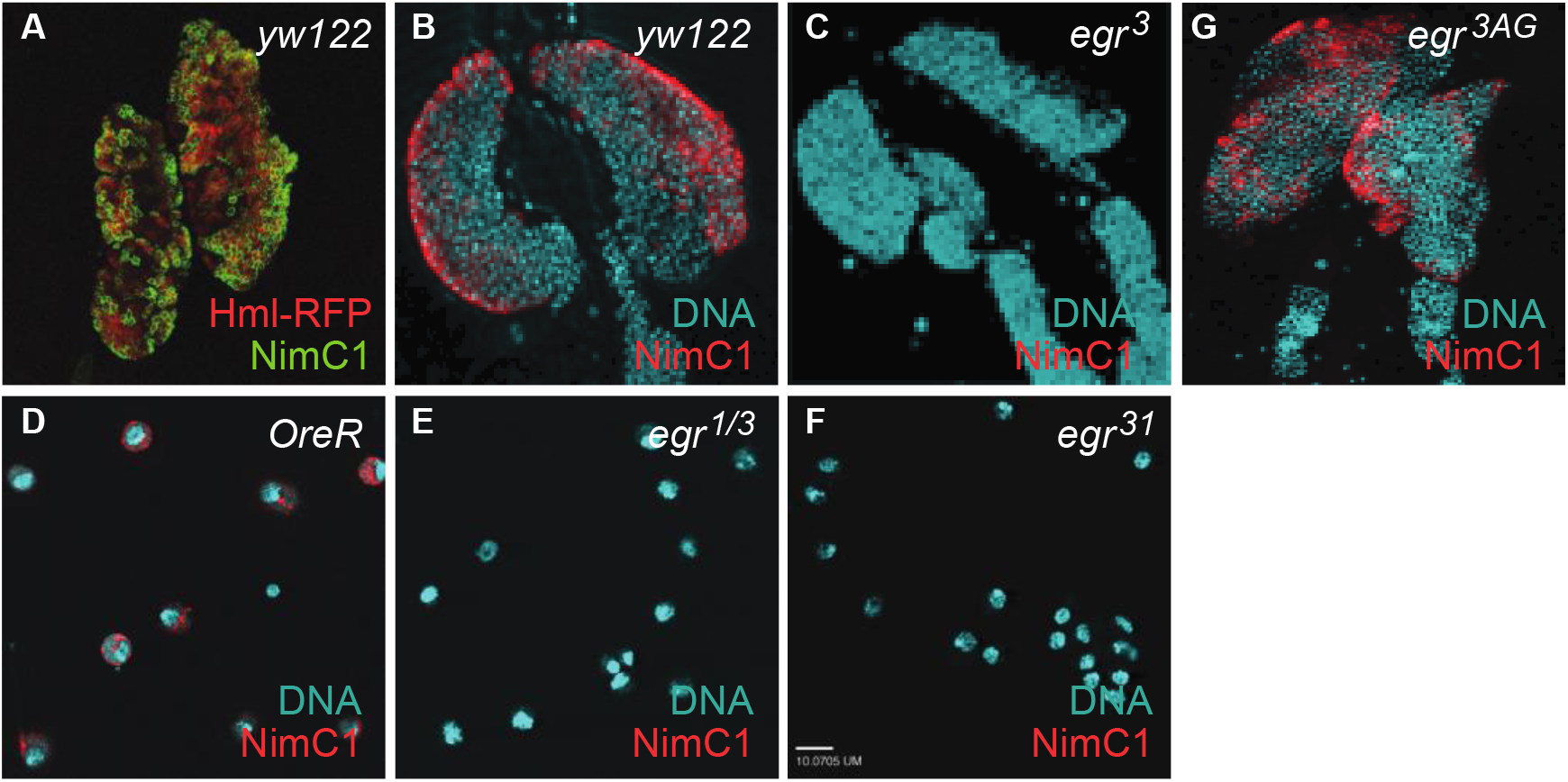
Lymph glands from *egr^3^* mutant larvae and circulating hemocytes from *egr^1/3^* and *egr^31^* mutant larvae are negative for NimC1 staining. A. Hml-RFP (red) and NimC1 (green) are expressed in many hemocytes in lymph glands from control, *yw122* larvae. B. NimC1 (red) staining in the primary lobes of lymph glands from control larvae. C. Lymph glands from *egr^3^* mutant larvae have no NimC1-positive hemocytes. D. Circulating hemocytes from the hemolymph of OreR control larvae stain positively for NimC1 (red). E-F. Circulating hemocytes from *egr^1/3^* transheterozygous larvae (E) and from *egr^31^* mutant larvae (F) lack positivity for NimC1. G. NimC1 staining (red) in the *egr^3AG^* mutant lymph glands, in which the *NimC1* locus was restored to WT.

Honti and colleagues reported that several *Drosophila* strains that were negative for NimC1 staining carried mutations in the *NimC1* gene, which they postulated were scars of mobile element mobilization (Honti *et al.*). Genomic sequencing of these P1-negative strains identified two independent micro-deletions in the *NimC1* gene, including a 6 bp deletion between nucleotides 2264 to 2270 (Honti *et al.*). Another deletion of 355 bp was found between nucleotides 1582 to 1937, accompanied by a 5 bp insertion. Together, Honti et al found that the 355 bp deletion and the 5 bp insertion generated a frameshift mutation in both the *NimC1 RA* and *RB* transcripts, resulting in new sequences and a premature stop codon (Honti *et al.*). The alterations were predicted to give rise to a truncated NimC1 protein that lacks the intracellular and transmembrane domains and four extracellular NIM repeats, which would account for its absence on the plasma membrane of hemocytes (Honti *et al.*).

To determine whether the *egr^1^* and *egr^3^* mutants carried mutations at the *NimC1* locus, we carried out genomic sequencing of a 1254 bp region that encompasses most of the *NimC1* open reading frame (**Fig. 3**). Our data shows that both *egr^1^* and *egr^3^* contain identical microdeletions and insertions within the *NimC1* gene, consistent with their common parental origin. Each mutant strain has the same 355 bp deletion, 5 bp micro-insertion, and 6 bp micro-deletion described by Honti *et al* at residues 2264-2270 (**Fig. 3B, C)**. In addition, using primers flanking the larger, 355 bp deletion in PCR reactions, we found that both the *Regg1^GS9830^* and *egr^31^* strains carried similar lesions. Since these *egr* mutants were both NimC1 negative (**Fig. 2E, F**), they very likely also carry the premature stop codon generated by the 355 bp deletion and 5 bp insertion. Altogether, these results suggest that these *NimC1* polymorphisms were present in the parental strain (**Fig. 3D**).

**Fig. 3.**
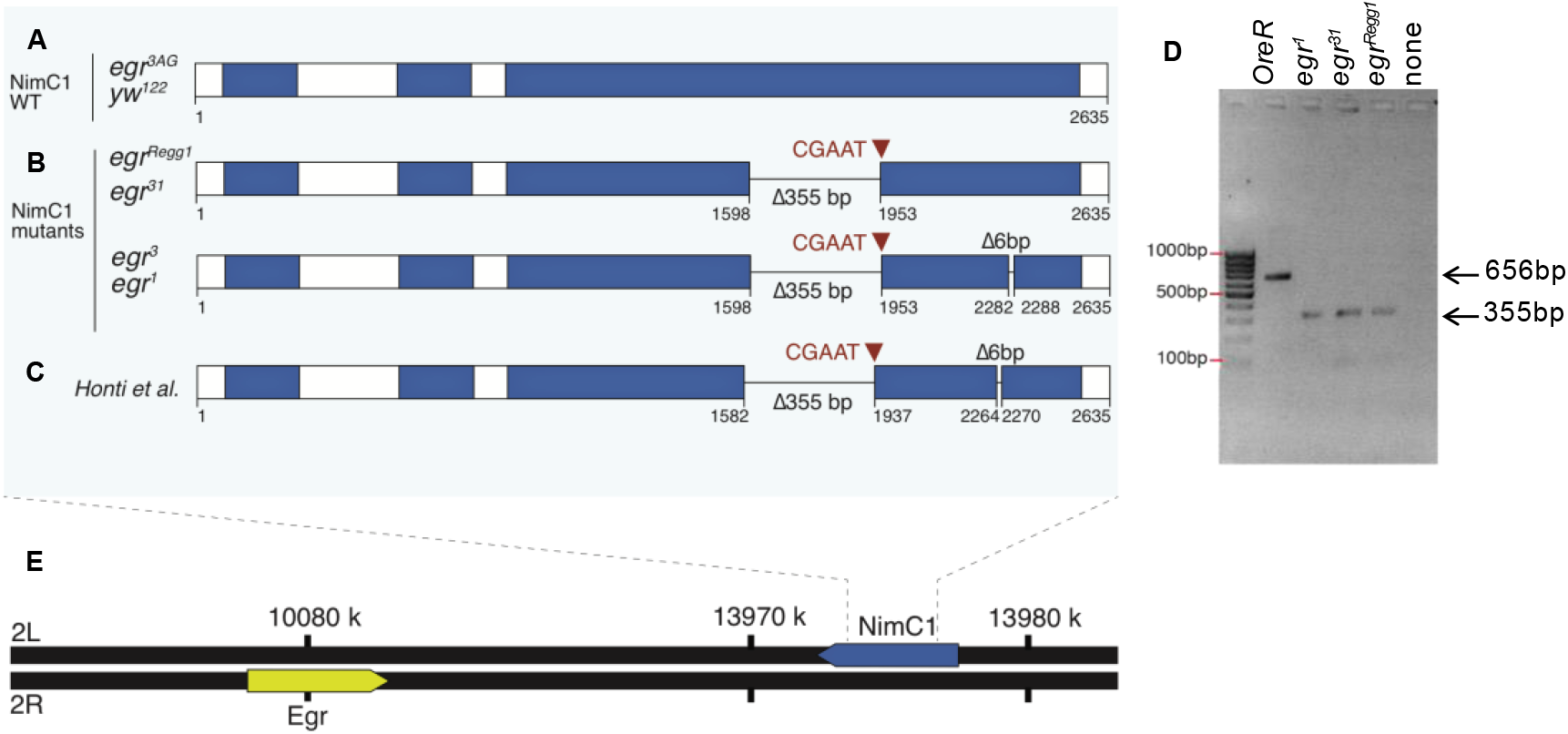
Summary of mutations in the *NimC1* locus in various *egr* mutants. A. Schematic representation of the *NimC1* locus, located at on the left arm of Chr. 2 (2L in D). White blocks represent introns and 5’ and 3’ untranslated regions. The sequence of the locus from *yw^122^* flies, used as a WT strain, and from the outcrossed *egr^3AG^* strain, is identical to the reference (*D. melanogaster* version r5.23). PCR genotyping suggests that the *OreR* WT strain is also wild type at the *NimC1* locus. Numbering is as in https://flybase.org/decoratedfasta/FBgn0259896. B. Representation of the *NimC1* locus in the *egr^1^* and *egr^3^* strains. Two deletions, of 355 bp and 6bp, and an insertion of 5 bp, are shown. PCR genotyping in *egr^31^* and the parental strain, *egŕ^Regg1^* indicates that they also carry the 355 bp deletion (D), and likely also the 5 bp insertion and 6 bp deletion. C. Representation of the *NimC1* locus from Honti *et al*, 2013. Note that the locus numbering is slightly different than in B, presumably due to an earlier genome annotation. The mutations are identical to those found in *egr^1^* and *egr^3^* and similar to *egr^31^* and the parental line, *egr^Regg1^* (B). D. Results of PCR genotyping of *NimC1* in *egr* mutants using primers flanking the 355 bp deletion and 5 bp insertion between nucleotides 1582 to 1937 (Honti *et al* genome annotation). E. Representation of Chr. 2, showing the *NimC1* locus on 2L, and the *egr* locus on the right arm of Chr. 2 (2R).

To restore the wild-type NimC1 locus to the *egr* mutants, we outcrossed both *egr^1^* and *egr^3^* to the *OreR* wild-type strain and isolated recombinants with the WT *NimC1* locus and either the *egr^1^* or *egr^3^* mutation (see Methods). We then sequenced the *NimC1* locus in these outcrossed *egr* alleles (hereafter called *egr^1AG^* and *egr^3AG^*) to verify that the recombination removed the mutant sequences. Both the *egr^1AG^* and *egr^3AG^* strains lacked the deletions and micro-insertions that characterized the original *egr^1^* and *egr^3^* alleles (**Fig. 3A**). Consistent with the loss of the deletions, hemocytes from the *egr^1AG^* and *egr^3AG^* mutants regained NimC1 positivity (**Fig. 2G** and data not shown).

Our sequencing data thus confirm that the *egr^1^* and *egr^3^* mutants also have mutations at the *NimC1* locus. Since the large deletion and micro-insertion in exon 3 of *NimC1* also exists in both the original parental line, *egr^Regg1^* and in *egr^31^*, a precise excision of the *Regg1^GS9830^* P-element, it is highly likely that the *NimC1* mutations in each of these *egr* alleles are derived from the parental strain. These *NimC1* mutations are recessive (Honti *et al.*), and their presence on each *egr* mutant chromosome in our experiments might explain the transiently persistent dying cells; perhaps they also account for the infection susceptibility found previously in *egr^3^* mutants (Schneider *et al.*). Consistent with our sequencing results, the genetic backgrounds of the *egr^1^* and *egr^3^* alleles were previously noticed to harbor anomalies that led to *egr*-independent susceptibility to infection by Gram-positive bacteria (Narasimamurthy *et al*. 2009). Complete deletion of *NimC1* has been reported to prevent phagocytosis of latex beads or yeast zymosan particles by plasmatocytes (Melcarne *et al.*), but whether and how phagocytosis of dying cells may be impaired by the *NimC1* mutations we found here remains to be determined. NIM repeats are thought to mediate protein-protein interactions and clustering of receptors are proposed to be key in phagocytic removal of apoptotic cells (Shklyar *et al.*). If the truncated mutant NimC1 proteins are aberrantly secreted into the hemolymph, as predicted (Honti *et al.*), they could interfere with critical NIM interactions. As the *egr^1^ and egr^3^* alleles have been used in numerous studies of immunity and cell death, it may be worthwhile to re-evaluate some of the phenotypes obtained with these alleles.

## Methods

### Fly strains and husbandry

Flies were raised at 25°C on standard cornmeal-molasses food supplemented with fresh dry yeast. The following strains were used: *egr^1^, egr^3^* and *egr^Regg1^* (Igaki *et al.), egr^31^* (gift of H. Kanda)*, egr^3AG^, egr^1AG^*(generated in this work)*, UAS-eg^IR^* (gift of M. Miura), *hml-RFP/CyO* (gift of K. Brückner)*, yw hsflp^122^* (gift of G. Struhl), *act>y^STOP^>Gal4* and *tub>Myc^STOP^>Gal4* (de la Cova *et al*.), *OregonR* (from Bloomington Drosophila Stock Center).

### Cell death assays

Eggs from appropriate crosses were collected on yeasted grape plates for 2-4 hours and allowed to develop at 25 °C in a humid chamber for 24 hours. At hatching, larvae were transferred to food vials supplemented with fresh yeast paste at densities of <50 larvae/vial to prevent crowding. To generate ‘loser’ clones in a competitive context, a *tub>Myc^STOP^>Gal4* cassette (> represents a FLP-recognition target (FRT) site) was used to excise the *>Myc^STOP^* cassette and generate *tub>Gal4, UAS-GFP* expressing “loser” cells in WT and in *egr^3^* mutants. Clones were allowed to grow in wing discs for 24 hours, as described (de la Cova *et al*.; Meyer *et al*.; Alpar *et al.*). FLP recombinase, under heat shock (HS) control, was activated by HS of larvae at 37°C for 10 minutes, at 48 hours after egg laying (AEL). Post-HS, larvae were allowed to grow at 25°C for 24 or 48 hours. To generate Myc-expressing clones, *act>y^STOP^>Gal4* was used to generate *act>Gal* clones that expressed UAS-GFP and UAS-Myc in WT and in *egr^3^* mutants. *act>Gal4* clones were induced in larvae with a HS at 37°C for 6 min. Wing discs were dissected from larvae at 48 hours after clone induction (ACI). A detailed protocol is available upon request.

### Larval dissection and imaging

Wing imaginal discs were dissected from third instar larvae as indicated above, and fixed in 4% paraformaldehyde in phosphate-buffered saline (PF-PBS) for 20 min at room temperature and washed with 0.01% Tween-20 in PBS (PBTw). Larval carcasses were stained with Rabbit anti-Cleaved Caspase-3 (Cas-3) at 1:100 (Cell Signaling). Secondary antibodies were purchased from Jackson Immunoresearch. Hoechst 33258 was used to stain DNA. Lymph glands were stained with plasmatocyte-specific P1 antibodies (anti-NimC; 1:100) (Kurucz *et al*. 2007) as described below for hemocytes. Wing discs and lymph glands were mounted in VectaShield Antifade on glass slides. Images were acquired with a Zeiss Axiophot microscope with Apotome and processed using ImageJ and Adobe Photoshop.

### Hemocyte immunohistochemistry and image processing

Hemocytes were collected from 10-20 experimental larvae, by bleeding from a small tear in the posterior cuticle into a 10-fold volume of PBS. Cells were then transferred to a coverslip and allowed to settle for 30 minutes at room temperature in a humidity chamber. All subsequent steps were performed directly on the coverslip. Cells were fixed in 4% PFA in PBS for 7 minutes at room temperature, washed 3 times in PBS, permeablized for 5 minutes in 1% triton in PBS (PBT), blocked for 5 minutes in 10% normal goat serum (NGS) in PBT and then incubated with primary antibody in 10% NGS in PBT either overnight at 4 °C or for 30 minutes at room temperature. Washes were carried out in PBT and secondary incubation was performed for 30 minutes at room temperature in 10% NGS in PBT. Cells were then washed 2 times in PBT, followed by 2 washes in PBS. A final 5 minute incubation with DAPI in PBS was performed. Coverslips were mounted in Slowfade Gold Antifade (Molecular Probes). Primary antibodies used were plasmatocyte-specific anti-P1 (1:100) (Kurucz *et al*. 2007). Secondary antibodies were Alexa Fluor antibodies (1:500, Invitrogen). Images were collected on a Zeiss Axio Imager M1 and were processed using ImageJ and Adobe Photoshop.

### Genomic sequencing of *egr* alleles

Genomic DNA was isolated from homozygous adult female flies. 100 ng of DNA was amplified using the primer sets as described in Honti et al. 2013. The 355bp deletion was found in the fragment amplified by P11189fw (CGCAGGAGCCTACGATAATC) and P11189rev (AAGGAATGTGGACACCATAG). The 34bp insertion was detected by the primers P1utan600fw (AACTGGATCGTCTAACAAGT) and P1utan300rev (GGATTGATTAACCACACAGA). The fragments were cloned into a pCR™4-TOPO-TA vector for sequencing using the common sequencing primers M13. Primers are listed in Table 1.

**Table 1.**
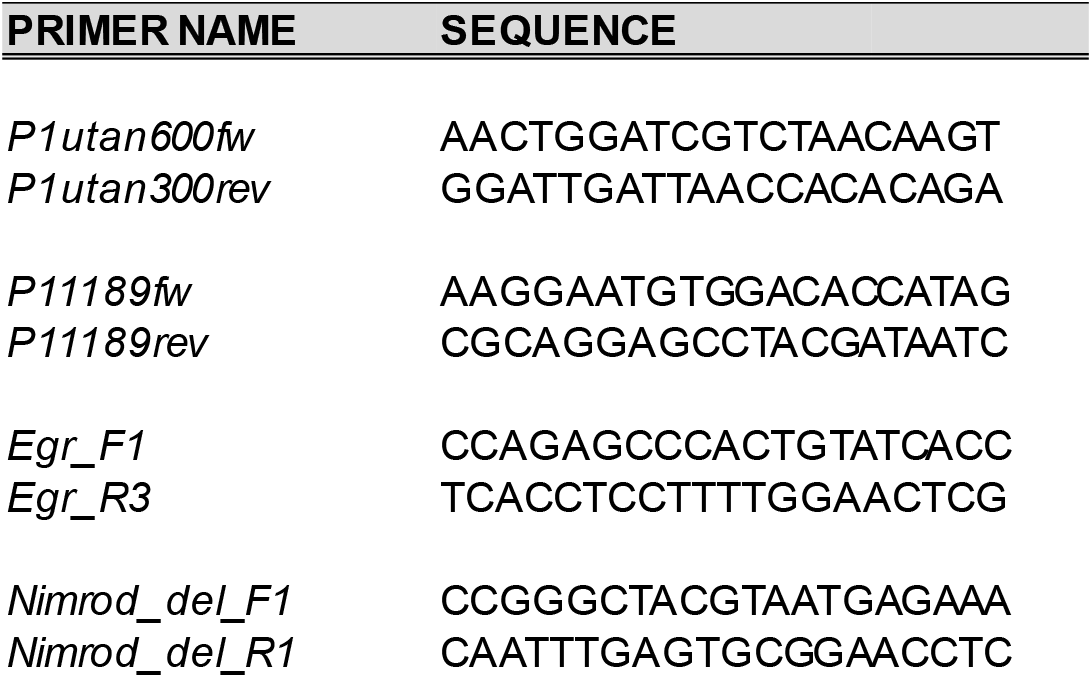
Primers used in this study

### Outcrossing and genotyping of *egr* alleles

*egr^1^ and egr^3^* alleles (hereafter *egr*) were treated identically. *egr* mutant virgin females were crossed to *OregonR (OreR*) males and the resulting F1 heterozygous *egr/+* virgin females were backcrossed to *OreR* males. Ten F2 males (either *egr/+* or +/+) were singly crossed to virgin *OreR* females, sacrificed and used for single-fly PCR (Gloor *et al.*) to identify males carrying the *egr* alleles, but no longer carrying the 355bp deletion in *NimC1*. A single *egr*-positive, *NimC1*-negative line for each allele was carried forward by crossing F3 virgin females back to *OreR* males. This process was repeated four times, after which *egr* alleles were re-isolated by crossing to *+; Sco/CyO actin-GFP*, a chromosome II marker/balancer strain similarly crossed into an *OreR* background. These alleles were named *egr^1AG^* and *egr^3AG^*. Genotyping of *egr^1^* and *egr^3^* used primers flanking the reported deletion in each strain: Egr_F1 (CCAGAGCCCACTGTATCACC) and Egr_R3 (TCACCTCCTTTTGGAACTCG) amplify a ~1500bp and ~2000bp fragment in *egr^1^* and *egr^3^*, respectively. Genotyping of *NimC1* used primers flanking the 355bp deletion and 5bp insertion between nucleotides 1582 to 1937 (Honti genome annotation). Nimrod_del_F1 (CCGGGCTACGTAATGAGAAA) and Nimrod_del_R1 (CAATTTGAGTGCGGAACCTC) amplify a 656bp fragment in WT animals and a ~300bp fragment in animals bearing the *NimC1* deletion. Primers are listed in Table 1.

## Acknowledgements

We are grateful to Chris Cary for technical assistance, and members of the Johnston lab for advice. We thank Masayuki Miura, Tatsushi Igaki, Hiroshi Kanda and Iswar Hariharan for providing fly stocks, and István Andó for the gift of anti-NimC1 antibodies. We are indebted to the Bloomington Drosophila Stock Center (BDSC) (NIH P40OD018537) and Flybase (Attrill *et al.*) for their services. Funding for this work was provided by the NIH RO1GM078464 and NCI RO1CA192838 (to LAJ) and the NSF (Graduate Research Fellowship to ARG). The *egr^3AG^* and *egr^1AG^* alleles, both WT for *NimC1*, were isolated by ARG in the lab of Iswar Hariharan and will be deposited at the BDSC for general use.

## References

Abrams, J. M., K. White, L. I. Fessler and H. Steller, 1993 Programmed cell death during Drosophila embryogenesis. Development 117: 29–43.

Alpar, L., C. Bergantinos and L. A. Johnston, 2018 Spatially Restricted Regulation of Spatzle/Toll Signaling during Cell Competition. Dev Cell 46: 706–719 e705.

Attrill, H., K. Falls, J. L. Goodman, G. H. Millburn, G. Antonazzo et al., 2016 FlyBase: establishing a Gene Group resource for Drosophila melanogaster. Nucleic Acids Res 44: D786–792.

Chung, Y. S., and C. Kocks, 2011 Recognition of pathogenic microbes by the Drosophila phagocytic pattern recognition receptor Eater. J Biol Chem 286: 26524–26532.

Clark, R. I., K. J. Woodcock, F. Geissmann, C. Trouillet and M. S. Dionne, 2011 Multiple TGF-beta superfamily signals modulate the adult Drosophila immune response. Curr Biol 21: 1672–1677.

de la Cova, C., M. Abril, P. Bellosta, P. Gallant and L. A. Johnston, 2004 Drosophila myc regulates organ size by inducing cell competition. Cell 117: 107–116.

Franc, N. C., P. Heitzler, R. A. Ezekowitz and K. White, 1999 Requirement for croquemort in phagocytosis of apoptotic cells in Drosophila. Science 284: 1991–1994.

Gloor, G. B., C. R. Preston, D. M. Johnson-Schlitz, N. A. Nassif, R. W. Phillis et al., 1993 Type I repressors of P element mobility. Genetics 135: 81–95.

Honti, V., G. Cinege, G. Csordas, E. Kurucz, J. Zsamboki et al., 2013 Variation of NimC1 expression in Drosophila stocks and transgenic strains. Fly (Austin) 7: 263–266.

Igaki, T., H. Kanda, Y. Yamamoto-Goto, H. Kanuka, E. Kuranaga et al., 2002 Eiger, a TNF superfamily ligand that triggers the Drosophila JNK pathway. EMBO J 21: 3009–3018.

Kocks, C., J. H. Cho, N. Nehme, J. Ulvila, A. M. Pearson et al., 2005 Eater, a transmembrane protein mediating phagocytosis of bacterial pathogens in Drosophila. Cell 123: 335–346.

Kurucz, E., R. Markus, J. Zsamboki, K. Folkl-Medzihradszky, Z. Darula et al., 2007 Nimrod, a putative phagocytosis receptor with EGF repeats in Drosophila plasmatocytes. Curr Biol 17: 649–654.

Manaka, J., T. Kuraishi, A. Shiratsuchi, Y. Nakai, H. Higashida et al., 2004 Draper-mediated and phosphatidylserine-independent phagocytosis of apoptotic cells by Drosophila hemocytes/macrophages. J Biol Chem 279: 48466–48476.

Melcarne, C., B. Lemaitre and E. Kurant, 2019a Phagocytosis in Drosophila: From molecules and cellular machinery to physiology. Insect Biochem Mol Biol 109: 1–12.

Melcarne, C., E. Ramond, J. Dudzic, A. J. Bretscher, E. Kurucz et al., 2019b Two Nimrod receptors, NimC1 and Eater, synergistically contribute to bacterial phagocytosis in Drosophila melanogaster. FEBS J 286: 2670–2691.

Meyer, S. N., M. Amoyel, C. Bergantinos, C. de la Cova, C. Schertel et al., 2014 An ancient defense system eliminates unfit cells from developing tissues during cell competition. Science 346: 1258236.

Milan, M., S. Campuzano and A. Garcia-Bellido, 1997 Developmental parameters of cell death in the wing disc of Drosophila. Proc Natl Acad Sci U S A 94: 5691–5696.

Moreno, E., M. Yan and K. Basler, 2002 Evolution of TNF signaling mechanisms: JNK-dependent apoptosis triggered by Eiger, the Drosophila homolog of the TNF superfamily. Curr Biol 12: 1263–1268.

Narasimamurthy, R., P. Geuking, K. Ingold, L. Willen, P. Schneider et al., 2009 Structure-function analysis of Eiger, the Drosophila TNF homolog. Cell Res 19: 392–394.

Schneider, D. S., J. S. Ayres, S. M. Brandt, A. Costa, M. S. Dionne et al., 2007 Drosophila eiger mutants are sensitive to extracellular pathogens. PLoS Pathog 3: e41.

Shklyar, B., F. Levy-Adam, K. Mishnaevski and E. Kurant, 2013 Caspase activity is required for engulfment of apoptotic cells. Mol Cell Biol 33: 3191–3201.

Somogyi, K., B. Sipos, Z. Penzes, E. Kurucz, J. Zsamboki et al., 2008 Evolution of genes and repeats in the Nimrod superfamily. Mol Biol Evol 25: 2337–2347.

Sonnenfeld, M. J., and J. R. Jacobs, 1995 Macrophages and glia participate in the removal of apoptotic neurons from the Drosophila embryonic nervous system. J Comp Neurol 359: 644–652.

